# SID-4/NCK-1 is important for dsRNA import in *Caenorhabditis elegans*

**DOI:** 10.1101/702019

**Authors:** Sonya Bhatia, Craig P. Hunter

## Abstract

RNA interference (RNAi) is sequence-specific gene silencing triggered by double-stranded (ds)RNA. When dsRNA is expressed or introduced into one cell and is transported to and initiates RNAi in other cells, it is called systemic RNAi. Systemic RNAi is very efficient in *C. elegans* and genetic screens for **s**ystemic RNA**i d**efective (Sid) mutants have identified RNA transporters (SID-1, SID-2 and SID-5) and a signaling protein (SID-3). Here we report that SID-4 is *nck-1*, a *C. elegans* NCK-like adaptor protein. *sid-4* null mutations cause a weak, dose-sensitive, systemic RNAi defect and can be effectively rescued by SID-4 expression in target tissues only, implying a role in dsRNA import. SID-4 and SID-3 (ACK-1 kinase) homologs interact in mammals and insects, suggesting they may function in a common signaling pathway, however, a *sid-3; sid-4* double mutants showed additive resistance to RNAi, suggesting that these proteins likely interact with other signaling pathways as well. A bioinformatic screen coupled to RNAi sensitivity tests identified 23 additional signaling components with weak RNAi defective phenotypes. These observations suggest that environmental conditions may modulate systemic RNAi efficacy, and indeed, *sid-3* and *sid-4* are required for growth temperature effects on systemic RNAi silencing efficiency.

## Introduction

RNAi is sequence-specific gene silencing triggered by natural or experimentally introduced double-stranded RNA (dsRNA) (Fire *et al.* 1998). In many animals, including *C. elegans*, experimentally introduced dsRNA is mobile (Jose and Hunter 2007). This dsRNA mobility allows dsRNA introduced by localized injection, transgenic expression, and in some animals by ingestion, to produce a whole-animal systemic silencing response. We previously reported the results of a visual screen for **s**ystemic RNA**i d**efective (Sid) mutants that has led to the identification of dsRNA transport and signal transduction proteins (Winston *et al.* 2002; Winston *et al.* 2007; Hinas *et al.* 2012; Jose *et al.* 2012).

Characterization of the *sid* genes has so far identified four SID proteins. SID-1 is a dsRNA channel protein that selectively transports long dsRNA into cells (Feinberg and Hunter 2003; Shih and Hunter 2011). In *sid-1* mutants, systemic RNAi is undetectable, but expressed or injected dsRNA can cause robust autonomous RNAi (Winston *et al.* 2002). Genetic mosaic and tissue-specific rescue experiments demonstrate that SID-1 is required for import but not export of silencing information (presumably dsRNA) (Winston *et al.* 2002; Jose *et al.* 2009; Whangbo *et al.* 2017). SID-2 is an intestinally expressed transmembrane protein, present at the intestinal lumen, that selectively endocytoses ingested dsRNA (Winston *et al.* 2007; McEwan *et al.* 2012). While SID-2 is required only for feeding RNAi, SID-1 is also required for feeding RNAi thus, it is presumed that SID-1 releases endocytosed dsRNA into the cytoplasm to initiate RNAi. SID-3 is a broadly expressed ACK1 tyrosine kinase homolog (Jose et al. 2012). *sid-3* mutants are partially defective for import of silencing signals. The requirement of a signal transduction protein suggests that environmental or physiological conditions may regulate systemic RNAi. SID-5 is a small novel protein associated with late endosomes (Hinas *et al.* 2012). Like *sid-3* mutants, *sid-5* mutants are partially defective for RNAi but *sid-5* is also essential for initiation of parental RNAi (transgenerational silencing) (Wang and Hunter 2017). Here we report the identification and characterization of *sid-4*.

Several other activities are also required for systemic RNAi. RME-2 is an endocytosis receptor that, in the absence of SID-1, is required to transport dsRNA into oocytes to support parental RNAi (Wang and Hunter 2017). Similar to feeding RNAi, SID-1 activity is subsequently required in the embryo for RNAi silencing, likely to release endosome trapped dsRNA into the cytoplasm to initiate RNAi. In addition, mutations in *rde-10 −11, −12*, and *rrf-1*,which fail to produce abundant secondary siRNAs resulting in dose-dependent RNAi silencing defects (Yang *et al.* 2014), are also required for effective systemic RNAi. dsRNA introduced locally (injection or expression) in *rde-12* and *rrf-1* mutants results in RNAi silencing locally, but not systemically, indicating that limiting amounts of dsRNA are transported. Thus, systemic RNAi is likely a dose-sensitive process. Our analysis of *sid-4* provides additional evidence for this hypothesis.

## Materials and Methods

### Strains

Unless otherwise indicated, all strains were grown at 20°C on NGM plates seeded with OP50 as a food source (Brenner 1974). Strains used in this analysis are listed in (Table 2 and for the candidate SID-3-SID-4 interacting screen (Table S1.

### Whole genome sequencing

Genomic DNA was extracted from each of the four *sid-4* strains: HC259 *sid-4 (qt15)*; HC119 *sid-4 (qt17)*; HC158 *sid-4 (qt33)*; and HC160 and *sid-4(qt35)*. Samples were sheared to 250bp and were prepared according to NEB Ultra DNA library kit E7370S. Sample concentrations were quantified via qPCR (Kapa Kit KK4824) and sequenced (Illumina), recovering a total of 250 million reads with 80x coverage. Reads were aligned using bowtie2 and exon variants on linkage group X were called using Samtools. Coding sequence variants in only a single gene, *nck-1* were identified in all four strains.

### SID-4 isoform expression constructs

#### Feeding RNAi assays

L4-young adults were placed on RNAi food (Kamath and Ahringer 2003) at room temperature (unless otherwise specified) and in every case, F1 adult progeny were scored for sensitivity to RNAi. Bacteria containing the empty-vector L4440 was used as the control for all feeding RNAi experiments.

#### sid-4 import assay

HC1139 *sid-4(ok694); Ex159 sid-4::gfp* young adults were cultured on *bli-1* RNAi plates for three days. Non-Bli animals were then moved to a separate plate and analyzed under a fluorescent microscope for GFP mosaics. Blistered animals on the *bli-1* RNAi plates were counted. All resistant worms expressing GFP were scored by widefield microscopy for GFP expression in the hypodermis.

#### dsRNA injections

*pal-1* dsRNA was made by amplifying a 1.2 kb region from the *pal-1* plasmid (Ahringer library, Kamath and Ahringer 2003) using forward and reverse T7 primers (*pal-1*-F-T7 TAATACGACTCACTATAGGTCCCATTTTAGGCAGTGAGTTA; *pal-1-R* GTTGCCAGCTCGTTATTTTATTG; *pal-1*-F TCCCATTTTAGGCAGTGAGTTTA; *pal-1-R-T7* TAATACGACTCACTATAGGCTCGAGAAGAAAAAGAACGACAA). A T7-flash Ampliscribe kit was used to make single-stranded sense and anti-sense RNA strands which were annealed, quantified, diluted as necessary, and injected into either one or both gonad arms of *sid-4(ok694)*and N2 animals. Six hours after injection each recovered animal was singled to an OP50 plate. To score embryonic lethality, the adult was removed after 24 hours and laid eggs counted. Three days later the number of hatched progeny were counted.

### SID-3-SID-4-Interactor Screen

All known mammalian NCK and ACK physical interactors were obtained from BioGrid (build 3.4.140) and Human protein reference databases (release 9) and all mammalian network information for both was obtained from PathCards (version 4.4) (Keshava Prasad et al. 2009; Belinky et al. 2015; Chatr-Aryamontri et al. 2017). Cross referencing all NCK interactors against both ACK interactors and the network data and ACK interactors against the NCK network data identified 113 unique mammalian proteins. We used wormbase (WS262) to identify 116 *C. elegans* orthologs of these proteins as well as to identify viable mutations. 53 mutant strains were ordered from the *C. elegans* Genetic stock Center (cgc.umn.edu). To test each strain for RNAi defects young adult hermaphrodites were placed on *fkh-6* RNAi plates and *dpy-11* RNAi plates. Three days later the *fkh-6* RNAi plates were scored for presence of laid eggs and the *dpy-11* RNAi plates were scored for Dpy progeny.

### Microscopy

Figure 6 images were obtained with a Zeiss Axiovert 200m spinning disc confocal microscope (63X and 100X objectives) equipped with a Hamamatsu Orca-ER digital camera. All the other images were obtained with an Olympus SZX2-TR30PT fluorescent microscope (filter (GFP-470) equipped with a Hamamatsu digital camera. Image J was used to adjust contrast and brightness and Adobe illustrator was used to crop images.

### Data availability

Strains not available at the cgc are available upon request.

## Results

### *sid-4/nck-1* encodes the *C. elegans* ortholog of the mammalian non-catalytic region of tyrosine kinase adaptor protein

The Sid screen recovered four X-linked, non-complementing alleles that define the *sid-4* locus (Winston *et al.* 2002). Whole genome sequencing of these four strains identified protein altering nucleotide variants in ZK470.5 (*nck-1*) in each isolate: *sid-4(qt15)* [S338L], *sid-4(qt17)* [G162E], *sid-4(qt33)* [Y250-stop] and *sid-4(qt35)* [E132K] (Figure 1). For subsequent characterization we used the 1814 base pair (bp) deletion allele, *sid-4/nck-1 (ok694)* (Consortium 2012) (Figure 1). SID-4/NCK-1 is homologous to adaptor proteins that promote the formation of signaling protein complexes either at the plasma membrane or in the cytoplasm (Buday 1999). Interestingly, SID-3 is homologous to ACK1 (activated CDC42 kinase) (Jose *et al.* 2012) and mammalian and Drosophila ACK1 proteins physically interact with NCK proteins (Teo *et al.* 2001; Worby *et al.* 2002). A translational *nck-1 genomic::gfp* reporter is broadly expressed in embryos and adults in a manner similar to SID-3::GFP reporters, supporting the possibility that *sid-3* and *sid-4* may interact to support systemic RNAi (Mohamed and Chin-Sang 2011; Jose *et al.* 2012).

**Figure 1.**
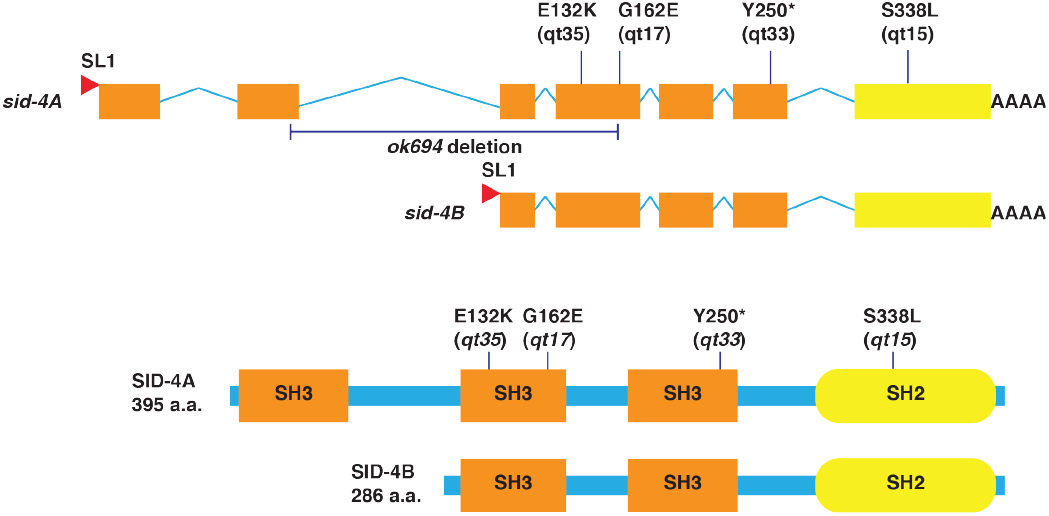
*sid-4* encodes NCK-1, a non-catalytic region of tyrosine kinase adaptor protein. Schematic of protein altering SNPs identified in ZK470.5 and their location in SID-4A and SID-4B (adapted from Mohamed et al. 2011). *sid-4A* and *sid-4B* are produced by independent promoters and trans-spliced to SL1.

### Both *sid-4* isoforms function to support systemic RNAi

The mammalian NCK homologs, *nck-1/ncka* and *nck-2*/nckβ, may function redundantly as well as have distinct roles in signal transduction (Chen *et al.* 1998; Li *et al.* 2001). *C. elegans sid-4/nck-1*, which is equally similar to both mammalian NCK homologs, encodes two isoforms: NCK-1A and NCK-1B (Figure 1). NCK-1A is the larger isoform and contains three SH3 domains followed by a single SH2 domain, while NCK-1B lacks the first SH3 domain (Figure 1). We obtained missense mutations in the SH2 domain and each of the SH3 domains common to both isoforms (Figure 1). The lack of a recovered mutation in the first SH3 domain fails to provide evidence of function for one or the other isoform. The two isoforms SID-4A and SID-4B are produced by two different *sid-4* promoters and transcription initiation sites (Figure 1) (Mohamed and Chin-Sang 2011). To test each isoform independently, we rescued the *sid-4(ok694)* with isoform specific cDNA constructs driven by their endogenous promoters.

To score RNAi efficacy in multiple tissues we monitored the expression of a *sur-5::gfp* transgene that expresses nuclear localized GFP in all somatic cells. The progeny of *sur-5::gfp* hermaphrodites grown on bacteria expressing *gfp* dsRNA are fully silenced, while roughly three-quarters of the progeny of *sid-4(ok694); sur-5::gfp* hermaphrodites grown on bacteria expressing *gfp* dsRNA are resistant or partially resistant to silencing (Figure 2). We then tested three independent SID-4A or SID-4B expressing extrachromosomal array lines for rescue of the silencing defect. GFP expression in non-intestinal cells was blind-scored in the progeny of animals placed on *gfp* RNAi food (Figure 2); the *sur-5::gfp* transgene line is not reliably expressed in the intestine in the presence of other extrachromosomal arrays (Jose *et al.* 2012). We found that both SID-4A and SID-4B similarly and partially rescued silencing in *sid-4; sur-5::gfp* strains (Figure 2). This result shows that expression of either SID-4 isoform is sufficient to produce SID-4 activity and, therefore that the first SH3 domain is not required for SID-4 to support systemic RNAi. However, the partial rescue suggest that the isoforms may function together for full activity. To test this, we injected both SID-4A and SID-4B cDNA constructs into the *sid-4(ok694); sur-5::gfp* strain, however all identified transgenic progeny were developmentally arrested. This suggest the two isoforms may function synergistically to affect growth or physiology, and perhaps systemic RNAi.

**Figure 2.**
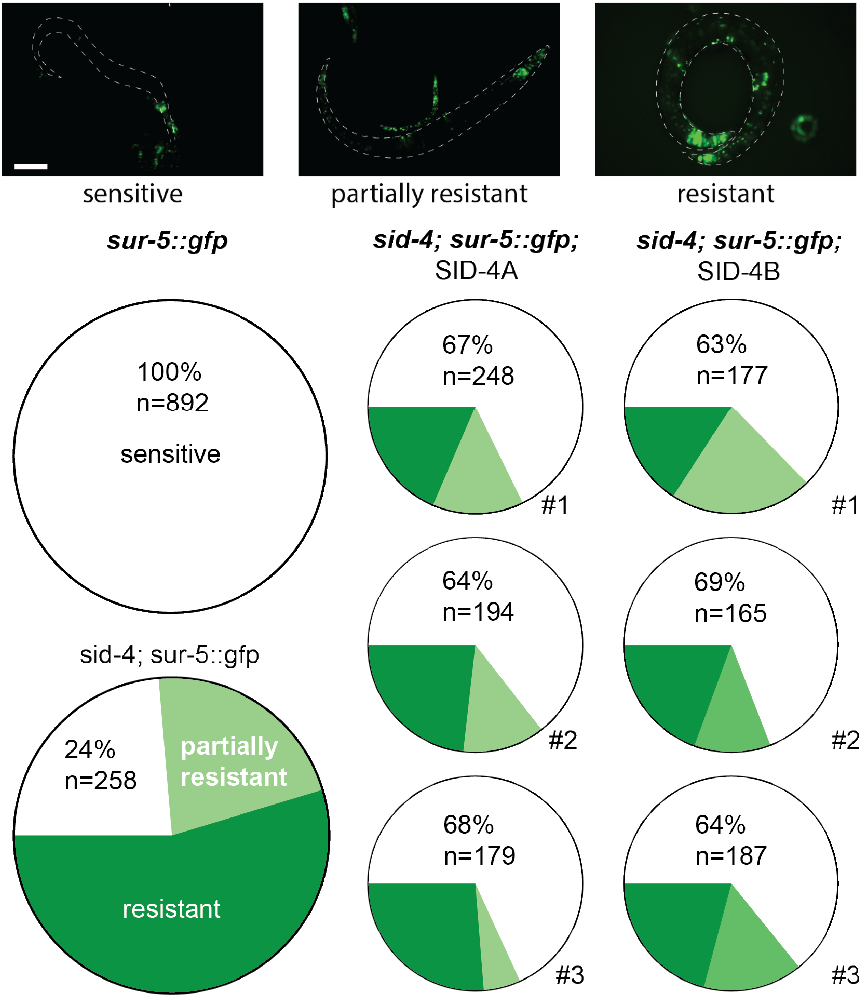
SID-4 isoform rescue of systemic RNAi. Representative images (top) of variable *sur-5*::GFP silencing. Quantification (bottom) of rescue of silencing in three independent lines for each SID-4 isoform. Scale bars: 0.1mm

### *sid-4* alleles have weak systemic RNAi defective phenotypes

The above analysis showed that *sid-4* mutants partially disable RNAi silencing. To directly compare *sid-4* systemic silencing defects to that of other Sid mutants, we crossed the deletion allele *sid-4(ok694)* into the transgenic background used for the screen. In this transgenic strain, pharyngeal expressed *gfp* dsRNA autonomously silences pharyngeal GFP and systemically silences GFP in anterior body wall muscle cells. When cultured on *gfp* dsRNA expressing bacteria, the ingested dsRNA silences GFP in all body wall muscle cells. A *sid-1* mutant in this transgenic background disrupts the silencing of the body wall muscle GFP from both pharyngeal expressed *gfp* dsRNA and ingested bacterial *gfp* dsRNA (Figure 3). In contrast, *sid-2* mutants disrupt only silencing in response to ingested dsRNA. Thus *sid-2* mutants in the transgenic background have the same silencing phenotype whether grown on normal bacteria or bacteria expressing *gfp* dsRNA (Figure 3). In contrast to *sid-1* and *sid-2* mutants, *sid-4(qt33)* and *sid-4(ok694)* show incomplete systemic RNAi defects (Figure 2A). Like *sid-1* and *sid-2*, neither allele disrupts pharyngeal silencing, confirming that the mutations do not noticeably disrupt RNAi. However, in contrast to *sid-1* and *sid-2*, many anterior body wall muscle cells are silenced when grown on bacteria expressing *gfp* dsRNA. While this indicates that *sid-4* is not required for uptake of ingested dsRNA, the lack of complete silencing may indicate a dose-dependent response to exported pharyngeal expressed or ingested *gfp* dsRNA. Detailed comparison of the extent of body-wall muscle GFP silencing shows no discernable difference between the two *sid-4* alleles (Figure 3).

**Figure 3.**
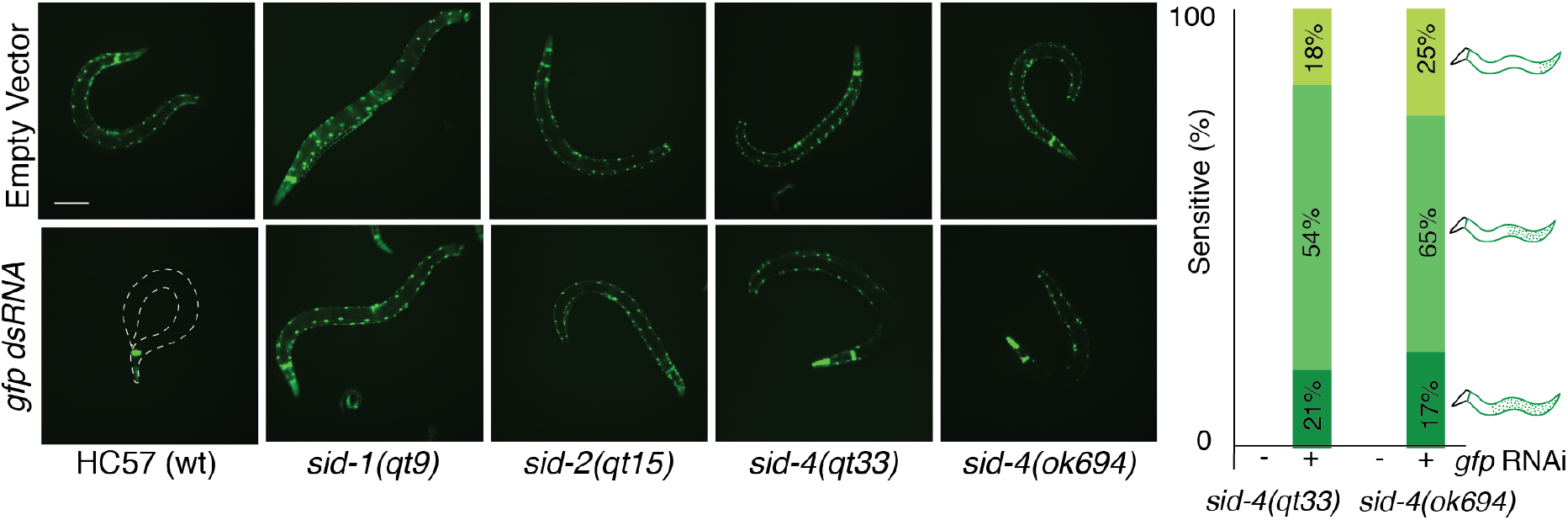
*sid-4* mutants are partially defective for RNAi. Adult F1 progeny of indicated *sid* mutant L4 hermaphrodites in the HC57 background (pharyngeal GFP; body wall muscle GFP; pharynx expressed *gfp* dsRNA) on bacteria expressing *gfp* dsRNA or empty vector (L4440 control). Scale bar 0.1mm. *sid-4 (ok694)* and *sid-4 (qt33)* progeny showed similar penetrance and expressivity of silencing defects as represented with accompanying schematics (n≥400). HC57 showed 100% sensitivity and *sid-1(qt9)* showed 0% sensitivity. Anticipated anterior muscle silencing in wt and *sid-2*animals on empty vector food is less than expected due to developmental stage imaged.

Weak *sid-1* alleles show gene-specific patterns of silencing that correlate with strong and weak RNAi foods (Whangbo *et al.* 2017). We tested both *sid-4* alleles on that same set of strong and weak RNAi foods. Wild-type worms were completely sensitive (100% silencing) to all these RNAi foods (Figure 4), while strong *sid-1* and *sid-2* mutant worms were completely resistant (0% silencing, not shown). *sid-4(qt33)* and *sid-4(ok694)* worms were completely or nearly completely resistant to *fkh-6* (gonad)*, unc-45* (muscle), and *bli-1* (hypodermis);completely or mostly sensitive to *act-5* (intestine) and *unc-22* (muscle); and partially resistant to *pos-1* (germline) and *dpy-11* (hypodermis). The discrepancy in sensitivity to *dpy-11* between *qt33* (nonsense) and *ok694* (deletion) indicates that the nonsense mutant *qt33* may have residual activity, perhaps reflecting read-through translation of the stop codon (Figure 4). The similarity in the pattern of RNAi sensitivity and resistance between strong *sid-4* alleles and weak *sid-1* alleles is consistent with *sid-4* mutations compromising systemic RNAi generally, rather than reflecting a tissue or dsRNA delivery specific effect.

Because *sid-3* is also a weak Sid mutant and mammalian and Drosophila SID-3 and SID-4 homologs have been shown to interact, we tested *sid-3(tm342)* on the same panel of RNAi foods (Figure 4). We observed a similar, but not identical pattern of resistance and sensitivity. These results are consistent with *sid-3* and *sid-4* acting in concert to promote systemic RNAi. In summary, these results indicate that the *sid-4* null phenotype is not a tissue-specific defect, but an incomplete RNAi defect.

**Figure 4.**
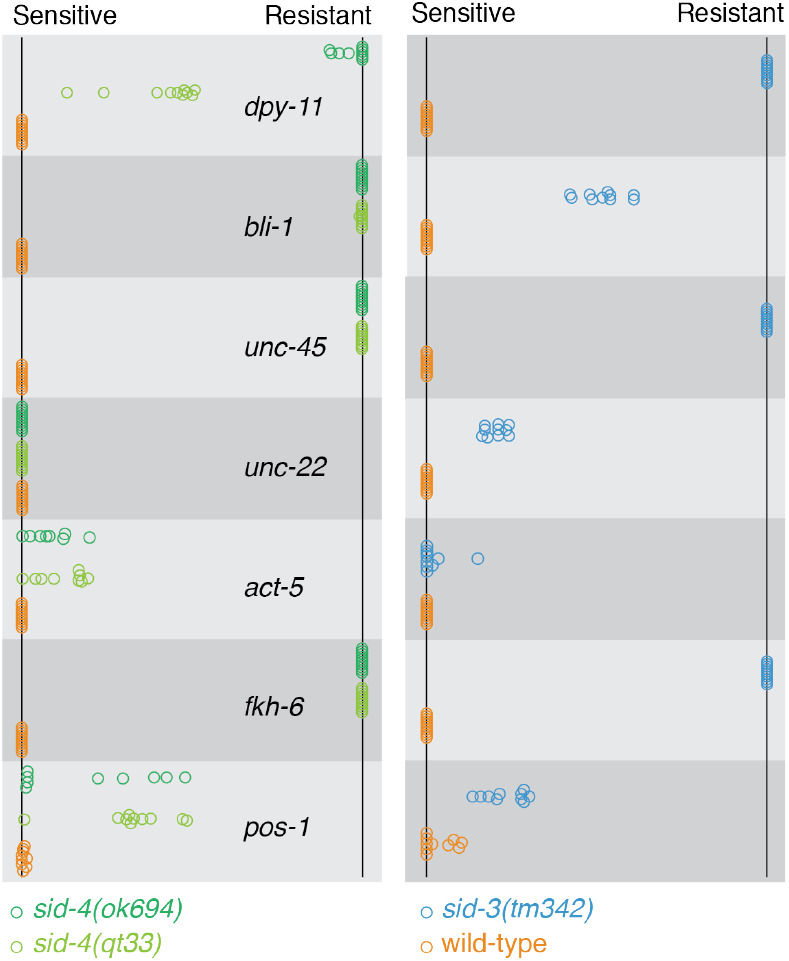
sid-4 and sid-3 have similar RNAi phenotypes. The adult progeny (F1) of L4 animals (F0) placed on RNAi foods were scored for RNAi sensitivity (fraction sensitive on each plate). Each circle represents the plate mean sensitivity for each F0 (n=10) with ≥150 F1’s for each F0. 100% sensitive (left) to 100% resistant (right).

### *sid-4* is a dose-dependent Sid mutant

*sid-4* mutant worms retain RNAi silencing activity in the pharynx and anterior body wall muscle cells (Figure 2A) indicating that *sid-4* mutations do not compromise RNAi silencing activity. However, weak RNAi defective (Rde) mutants that are suppressed by high levels of pharyngeal expressed dsRNA can produce wildtype silencing in the pharynx (Yang *et al.* 2014). To explicitly test *sid-4* mutants for a weak Rde phenotype we injected *pal-1* dsRNA directly into the syncytial germline of wild-type and *sid-4*mutant hermaphrodites and scored the frequency of the *pal-1*(RNAi) phenotype, embryonic lethality. Because the germline is syncytial, injections into the anterior or posterior gonad do not require dsRNA transport for effective silencing. If *sid-4* mutants are weak RNAi defective, then wild-type and *sid-4* mutant worms should show distinct embryonic lethality frequencies in response to decreasing injected dsRNA dose. We injected both gonad arms to eliminate the effect of gonad to gonad transfer. Furthermore, for each dsRNA concentration, we used a single needle and identical injection times to limit variability and enable direct comparison between strains. We found that both wild-type and *sid-4(ok694)* mutants showed proportionate changes in embryonic lethality when injected with changing concentrations of *pal-1* dsRNA (Figure 5). This indicates that *sid-4* mutants have a wild-type level of RNAi activity in the germline and are not weak Rde mutants.

**Figure 5.**
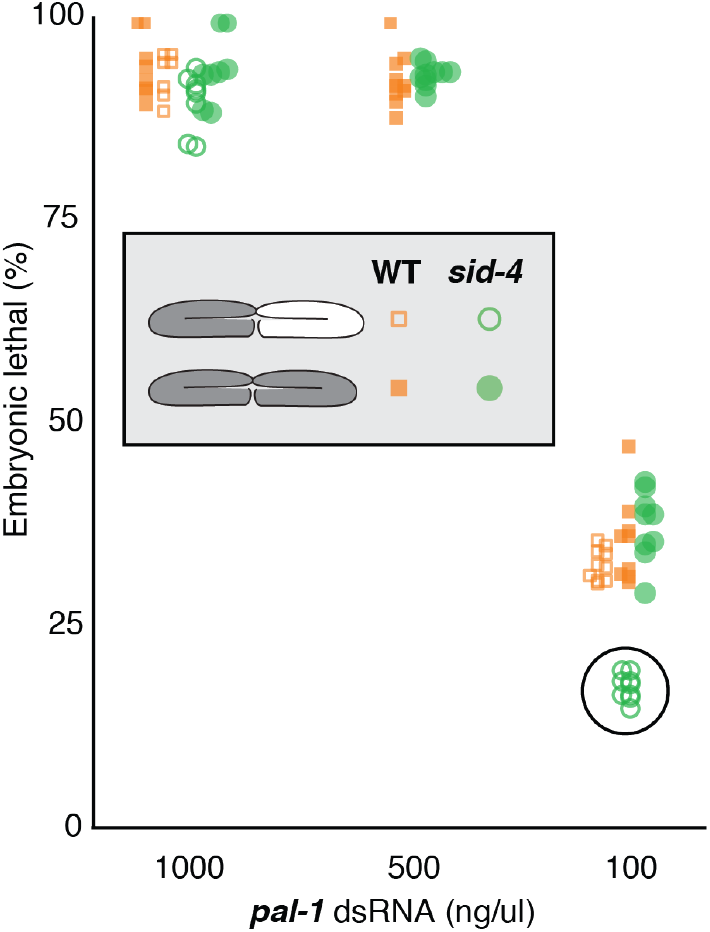
*sid-4* is a dose-dependent Sid. Wild-type and *sid-4(ok694)* young adults were injected with varying *pal-1* dsRNA concentrations into one or both gonad arms and then 6 hours post injection singled and allowed to lay eggs for 24 hours. The fraction of hatched eggs for each injected adult (n>10) was scored three days later.

We used the same assay to determine whether *sid-4* is required to transfer silencing information between gonad arms. In wild-type animals, injecting a single gonad arm can result in 100% embryonic lethality while in strong *sid-1* mutants only progeny from the injected gonad are affected (Winston *et al.* 2002; Wang and Hunter 2017). To test *sid-4(ok694)* mutants for systemic RNAi defects (Sid) we injected single gonad arms with either a high or low *pal-1* dsRNA dose. Consistent with effective systemic RNAi, there was no detectable difference in embryonic lethality among the progeny of wild-type worms injected with *pal-1* dsRNA into either a single or both gonad arms at either high or low dsRNA dose (Figure 5). Similarly, the frequency of embryonic lethality among the progeny of *sid-4(ok694)* animals injected with the high concentration of *pal-1* dsRNA into either a single or both gonad arms was nearly identical (Figure 5). However, the frequency of embryonic lethality among the progeny of *sid-4(ok694)* animals injected with the low concentration of *pal-1* dsRNA into single gonad arm was half that of animals injected into both gonad arms (Figure 5). This dose-dependent systemic RNAi result is consistent with inefficient transport of dsRNA between gonad arms, indicating that *sid-4* is a weak Sid mutant.

### *sid-4* is import defective

We used genetic mosaic analysis to determine whether *sid-4* is required in importing cells for efficient feeding RNAi. Specifically, we rescued *sid-4(ok694)* animals with a *sid-4::gfp* construct maintained as a mitotically unstable extrachromosomal array. About 82% of the progeny inherited the array and were GFP positive and formed full-body blisters when cultured on *bli-1* RNAi food. The remaining 18% of the progeny did not inherit the array, were mostly GFP negative and fully resistant to *bli-1* RNAi. *bli-1* is expressed in the large syncytial cell hyp7 (139 nuclei, 29 from embryonic cells that fuse to form hyp7 and 110 from postembryonic cells that fuse with hyp7) which produces the full body blister in *bli-1* RNAi treated animals (Chisholm and Hsiao 2012). Among 976 resistant worms we identified 74 with detectable GFP expression. If *sid-4* is required for import, then in these resistant animals we would expect all or most of the hyp7 nuclei to lack the *sid-4::gfp* array and fail to express detectable GFP in hyp7. Indeed, in none of these animals did we detect GFP expression in hyp7. It is notable that all of the hyp7 nuclei arise from descendants of three distinct early embryonic cells (ABarp, ABp, and C; Figure S1), thus to produce a *sid-4::gfp* mosaic lacking *sid-4::GFP* expression in most hyp7 nuclei would require two or three losses. For example (see figure S1B), the array may fail to segregate to AB at the first division and fail to segregate to P2 at the second division producing a mosaic animal that lacks *sid-4::gfp* in all hyp7 nuclei but expresses *sid-4::gfp* in the posterior pharynx, the gut, and anterior body wall muscle cells. 22 *bli-1* RNAi resistant animals were identified that had pharynx and gut GFP expression and lacked detectable GFP in hyp7 (Figure 6C). As a second example (Figure S1C), the array could be lost in P1, ABp, and ABar(pp), producing an animal that lacks *sid-4::gfp* in all hyp7 nuclei, but that expresses GFP in the anterior pharynx only. 52 animals were recovered that are consistent with this scenario (Figure 6D). It is important to note that by selecting for resistant animals that maintained some GFP, we greatly enriched for mosaic animals that failed to segregate the array at multiple cell divisions. In summary, the results of the *sid-4* genetic mosaic analysis indicate that *sid-4* is required in the target cell for effective RNAi.

**Figure 6.**
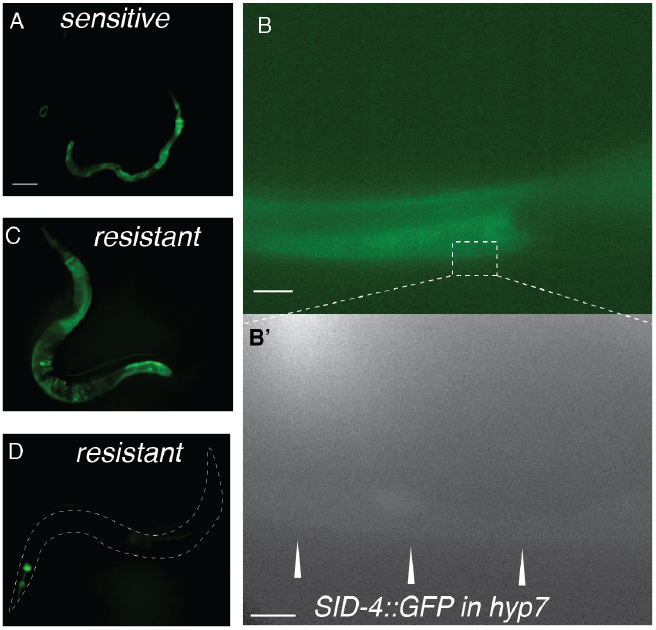
*sid-4* is required in the importing tissue. Progeny of *sid-4(ok694); [Ex:sid-4::gfp]* animals were scored for *bli-1* RNAi sensitivity and SID-4::GFP expression. A) A GFP positive *bli-1* RNAi sensitive animal. Scale bar: 0.1mm. B,B’) A confocal image of a *bli-1* RNAi sensitive animal showing GFP expression in hyp7. Scale bars: 25μm and 5μm respectively. C) Intestine and D) pharyngeal GFP expression in *bli-1* RNAi resistant animals.

### *sid-3* and *sid-4* genetically interact, but do not function in a simple linear pathway

Mammalian and *Drosophila* SID-3/ACK1 and SID-4/NCK-1 homologs interact (Teo *et al.* 2001; Worby *et al.* 2002), thus it is likely that they do so in *C. elegans* as well. Both *sid-4/nck-1* and *sid-3* fluorescent protein fusions constructs are broadly expressed cytoplasmic proteins that localize to intracellular membranes (Mohamed and Chin-Sang 2011; Jose *et al.* 2012). Furthermore, both *sid-3* and *sid-4* are weak Sid mutants required in importing cells for effective systemic RNAi (Jose *et al.* 2012) Figure 3, 4, 5, 6). If these genes function together as a kinase and exclusive adapter, then we expect the double mutant to resemble both single mutants. To test this, we constructed and tested a *sid-3(tm342); sid-4(ok694)* double mutant on a broad panel of RNAi foods (figure S2). For many of the foods the sensitivity of either single mutant was maximal or minimal, precluding interpretation of the double mutant sensitivity. But for two foods, *unc-22* (muscle) and *pos-1* (germline) the sensitivity of both single mutants was intermediate, and in both cases the double mutant was less sensitive (Figure 7A). This result shows that the effect of *sid-3* and *sid-4* are at least partially additive, indicating that in these two tissues these two signaling proteins do not function in a simple linear pathway, but likely interact independently with other signaling proteins to modulate systemic RNAi silencing.

**Figure 7.**
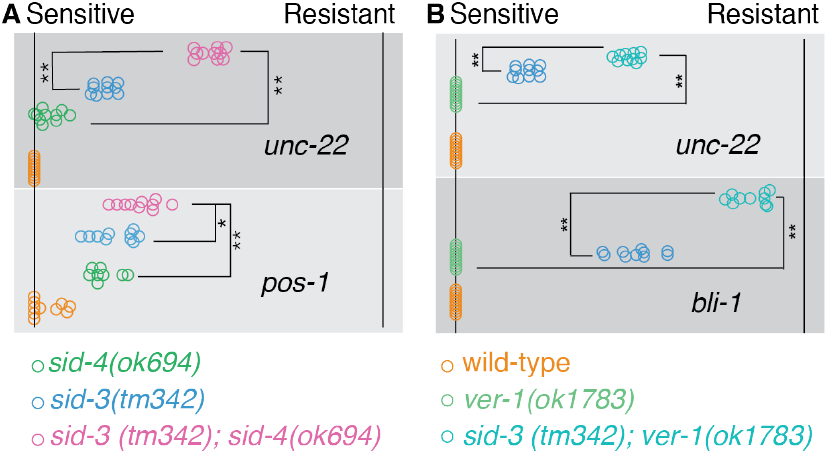
Double Sid weak mutants show enhanced RNAi resistance. The F_1_ adult progeny of F_0_ L4 animals placed on RNAi foods were scored for RNAi sensitivity (fraction sensitive on each plate). Each circle represents the plate mean sensitivity for each F_0_ (n=10) with ≥40 F_1_’s per F_0_. 100% sensitive (left) to 100% resistant (right). t-test p-values: * <0.01, ** <0.0005

### *C. elegans* homologs of mammalian NCK/ACK interacting proteins have weak RNAi phenotypes

The above double mutant analysis suggests that SID-3/4 interacting proteins may modulate systemic RNAi. To identify candidate *C. elegans sid-3* and *sid-4* interacting proteins we compiled a list 113 mammalian proteins that likely interact (directly or indirectly) with both a NCK and an ACK ortholog (see methods). Among 116 *C. elegans* orthologs of these proteins, 80 viable mutants have been described and we tested 52 by feeding RNAi for reduced sensitivity to *fkh-6* and *dpy-11* RNAi food (Table S1). 22 candidates were detectably resistant to *fkh-6* RNAi, seven of which were also partially resistant to *dpy-11* RNAi, and one strain was only partially resistant to *dpy-11* RNAi (Table 1). In addition, five of the 29 strains that were fully sensitive produced enhanced Dpy-11 phenotypes, possibly signifying enhanced RNAi.

**Table 1.** RNAi defective SID-3-SID-4 candidate interacting proteins

| Strain | Genotype* | *fkh-6* (RNAi)<br>eggs on<br>plate? (n=2) | *dpy-11* (RNAi)<br>% non-Dpy<br>adults (n=2) | Homology |
| --- | --- | --- | --- | --- |
| N2 | WT | none | 0 | --- |
| CB96 | *vab-2(e96) IV* | many | 44 | Ephrin Ligand |
| CZ375 | *vab-1(e856) II* | many | 0 | Ephrin Receptor |
| EM305 | *efn-4(bx80) IV* | many | 15 | Ephrin Ligand |
| RB942 | *cdc-42(ok825) II* | many/few | 19 | GTPase |
| VC610 | *ver-3(ok891) X* | many/few | 11 | Receptor Tyrosine Kinase |
| VC1263 | *ver-1(ok1738) III* | many | 0 | Receptor Tyrosine Kinase |
| RB2513 | *C26C6.6(ok3481) I* | many | 29 | Lim domain |
| MT12615 | *mys-1(n3681) V* | many | 0 | Histone Acetylase |
| VC664 | *ras-1(ok977) II* | few | 18 | GTPase |
| MT4434 | *ced-5(n1812) IV* | few | 15 | DOCK |
| CB3257 | *ced-2(e1752) IV* | few | 0 | SH2/SH3 adapter |
| CX51 | *dyn-1(ky51) X* | few | 0 | Dynactin |
| MT5267 | *soc-1(n1789) V* | few | 0 | Pleckstrin homology domain |
| RB1591 | *ddr-1(ok1956) X* | few | 0 | Tyr kinase |
| RB689 | *pak-1(ok488) X* | few | 0 | Ser/Thr kinase |
| NW1549 | *efn-2(ev658) IV;*<br>*efn-3(ev696) X* | few | 0 (n=1) | Ephrin Ligand |
| CZ414 | *vab-1(e699) II* | few | 0 | Ephrin Receptor |
| RB1267 | *ensh-1 (ok1349) X* | few | 0 | Fibrogenin like |
| RB1751 | *rga-5(ok2241) IV* | few | 0 | RhoGAP |
| RB759 | *akt-1(ok525) V* | few | 0 | Ser/Thr kinase |
| ZD500 | *hecw-1(ok1347) III* | few | 0 | E3 Ubiquitin Ligase |
| RB776 | *kin-32(ok166) I* | few | 0 | Tyr kinase |
| VC664 | *ras-1(ok977) II* | none | 18 | GTPase |

\*Some strains include non-relevant mutations, listed in full in Table S1.

Among the top candidates was the receptor tyrosine kinase VER-1 (Table 1), which is orthologous to the mammalian VEGF Receptor 1, an interactor of NCK1 (Igarashi et al 1998). In *C. elegans* VER-1 is expressed in the intestine and neuronal sheath cells (Procko et al 2012). To more thoroughly explore the RNAi sensitivity of the *ver-l(okl783)* deletion mutant, we scored the RNAi phenotypes of the progeny of adult animals placed on a panel of RNAi foods targeting a variety of tissues (Figure S3). The progeny of wild-type animals grown on *fkh-6* RNAi food fail to lay any eggs, and while *ver-l(okl783)* animals lay eggs (Table 1), however, this analysis showed that the proportion of egg-laying adults is small (Figure S3). Similarly, the penetrance of *dpy-11* RNAi resistance was low. These results suggest that *ver-1* is a very weak RNAi mutant. Because *sid-3, sid-4* double mutants show a stronger RNAi resistance phenotype than either single mutant, we constructed and tested *ver-l(okl783); sid-3(tm342)* double mutant on the same panel of RNAi foods (Figure S3). Consistent with VER-1 acting in the SID-4 pathway, we found that the double mutant was more resistant to feeding RNAi targeting *unc-22* and *bli-1* than either single mutant (Figure 7B).

In summary, 23 of 52 candidate SID-3/4 interacting protein mutants had detectable RNAi defects, supporting the hypothesis that additional signaling components and likely pathways modulate systemic RNAi. However, the low penetrance and expressivity of the RNAi defects combined with the high level of noise in the experimental assays makes it challenging to verify the significance of these interactions.

### SID-3/SID-4 are required for temperature-dependent RNAi silencing

Efficient systemic RNAi requires signaling proteins, indicating that physiological or environmental conditions may modulate systemic RNAi. We previously reported that temperature affects systemic RNAi; we noted more efficient silencing of body wall muscle GFP in response to exported pharyngeal *gfp* dsRNA at 20°C than at 25°C (Winston *et al.* 2002). To extend this analysis we determined whether temperature can affect the silencing of the endogenous gene *unc-22* in response to ingested *unc-22* dsRNA. The progeny of ten singled wild-type L4’s placed on *unc-22* RNAi food at 15°C, 20°C and 25°C were scored for *unc-22* twitching as adults. Consistent with the previous findings, two independent trials showed more efficient silencing at 15°C and less efficient silencing at 25°C (Figure 8). Thus, the temperature effect on systemic RNAi silencing is independent of the method of dsRNA delivery (expression vs ingestion) and target (transgene vs endogenous gene). We then repeated the experiment with *sid-3* and *sid-4* mutants (Figure 8). In contrast to wild-type, the progeny of both *sid-3* and *sid-4* worms showed less efficient silencing at all three growth temperatures. This experiment was then repeated in a separate large trial (Figure 8). These results indicate that *sid-3*and *sid-4* are required for the enhanced systemic RNAi silencing observed at lower temperature.

**Figure 8.**
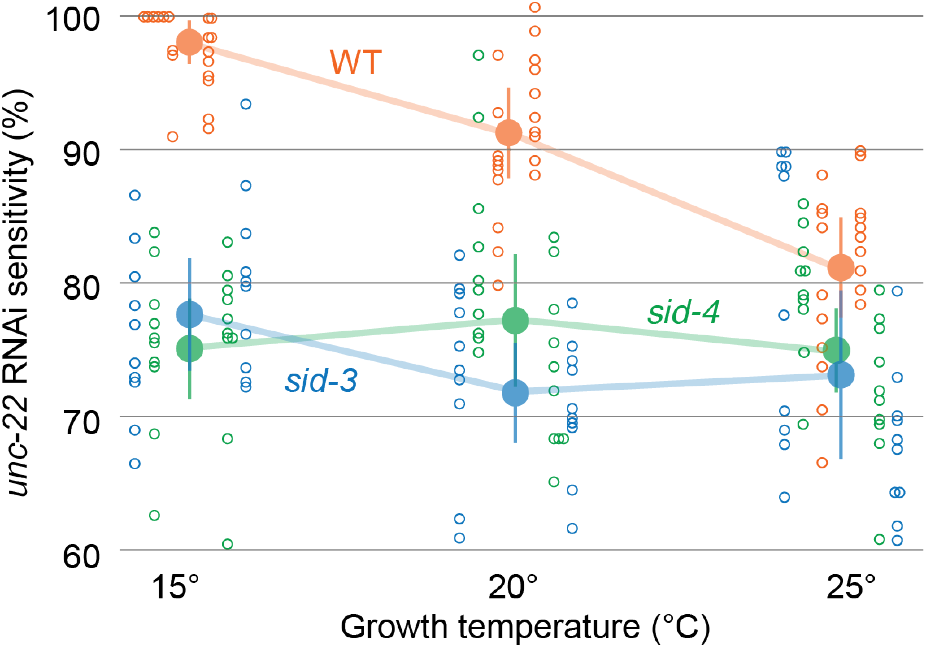
*sid-3* and *sid-3* temperaturedependent systemic RNAi. F_1_ progeny of F_0_ animals placed on *unc-22* food at the indicated temperature were scored in 3mM Levamisole for paralysis/twitching (sensitivity). Mean (large filled circle), 95% confidence interval (vertical line), and each F_0_ plate mean (small unfilled circle) are shown for each genotype-temperature. Plate means for two independent trials are shown to the left and right of each mean.

## Discussion

The identity of SID-4 as an NCK homolog complements the identity of SID-3 as an ACK1 homolog, one of the presumed SID-4/NCK-1 binding partners. Establishing that these proteins interact to regulate systemic RNAi in *C. elegans* and understanding what conditions regulate their activity will lead to even further insight into the mechanisms and functions of systemic RNAi. These two proteins have been implicated in a variety of presumably systemic RNAi-independent functions in *C. elegans*, including cell migration (Martynovsky *et al.* 2012) (Mohamed *et al.* 2012), virus entry (Jiang *et al.* 2017; Tanguy *et al.* 2017), and endocytic trafficking (Lazetic *et al.* 2018), and cell polarity (Goh *et al.* 2012). All these functions impinge on endocytosis.

Endocytosis has been implicated in feeding RNAi as well as SID-1-independent parental RNAi. In parental RNAi, the endocytosis receptor RME-2 is required for SID-1 independent transfer of extracellular dsRNA into oocytes (Wang and Hunter 2017) SID-1 is then subsequently required in the embryo for RNAi silencing, presumably to release the endocytosed dsRNA into the cytoplasm to initiate RNAi. In feeding RNAi, the SID-2 transmembrane protein, which is expressed in the intestine and localizes to the apical (lumenal) membrane, appears to act as a dsRNA specific endocytosis receptor (Winston *et al.* 2007; McEwan *et al.* 2012). SID-1 is also required for feeding RNAi induced silencing in intestinal cells, again, presumably to release endocytosed dsRNA into the cytoplasm to initiate RNAi. Furthermore, homologs of several of the candidate SID-3/SID-4 interacting proteins that have detectable Rde phenotypes have roles in or are implicated in endocytosis, including *dyn-1* (dynactin), *pak-1* (p21 activated kinase) and *cdc-42*. It should be noted that only viable alleles of candidate SID-3-SID-4 interacting proteins were tested and for many of these genes null alleles are inviable. Because endocytosis is essential for cell viability, our genetics screens will not recover strong Sid mutants disrupted for endocytosis.

The original large visual screen identified many strong alleles in *sid-1* and *sid-2* and relatively few weak alleles in *sid-3, sid-4* and *sid-5*. This bias likely reflects the relative ease of identifying worms with bright GFP in many cells in the strong mutants (*sid-1* and *sid-2* mutants) versus the variable and relatively sparse GFP expression in the weak mutants *(sid-3, sid-4*, and *sid-5*, as well as a partial loss-of-function *sid-1* alleles, (Whangbo *et al.* 2017). Our screen and analysis of candidate SID-3/4 interacting proteins extends this pattern, suggesting that most or all strong Sid mutants have been identified. However, our analyses also showed that *sid-3* and *sid-4* are dose-dependent SIDs and that combining two weak mutants resulted in stronger Sid phenotypes. Thus, sensitized screens that eliminate dsRNA dose effects on the phenotype or screens for enhancers or suppressors of weak alleles may identify additional genes and pathways important for systemic RNAi. Finally, analysis of systemic RNAi in *sid-3* and *sid-4* mutants identified temperature as an environmental factor that influences RNAi efficiency. Using temperature or similar environmental factors may provide additional avenues for identifying and understanding the contribution of these and other genes to systemic RNAi.

## Acknowledgements

We thank members of the Hunter lab, Alexander Weisman, Andrey Shubin, and Nicole Bush for comments of the manuscript. This work was supported by National Institutes of Health (NIH) grant GM089795. Some strains were provided by the Caenorhabditis Genetics Center, which is funded by NIH Office of Research Infrastructure Programs (P40 OD010440).

---

**Table S1.**
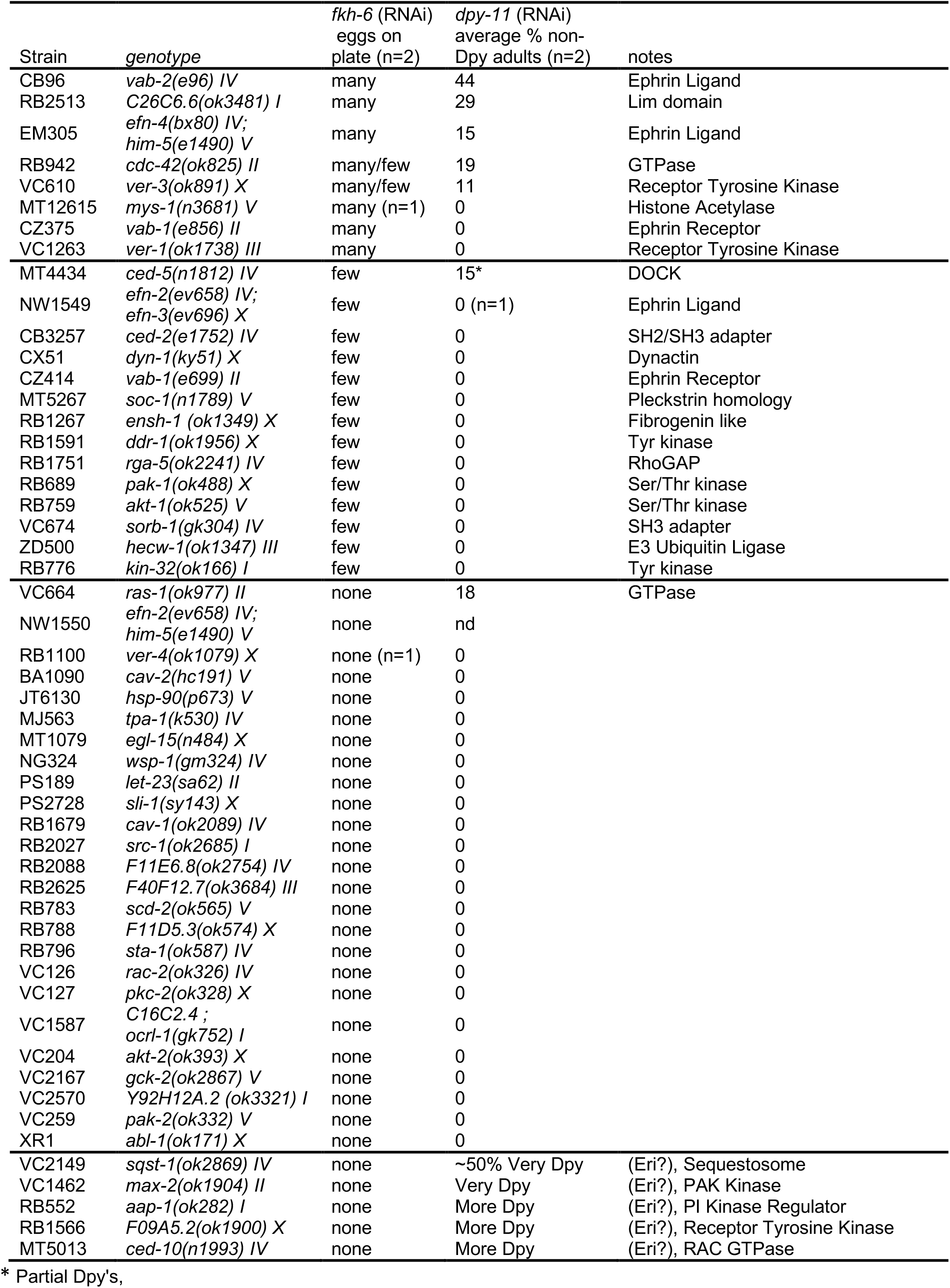

## Supplemental Figures

**Figure S1.**
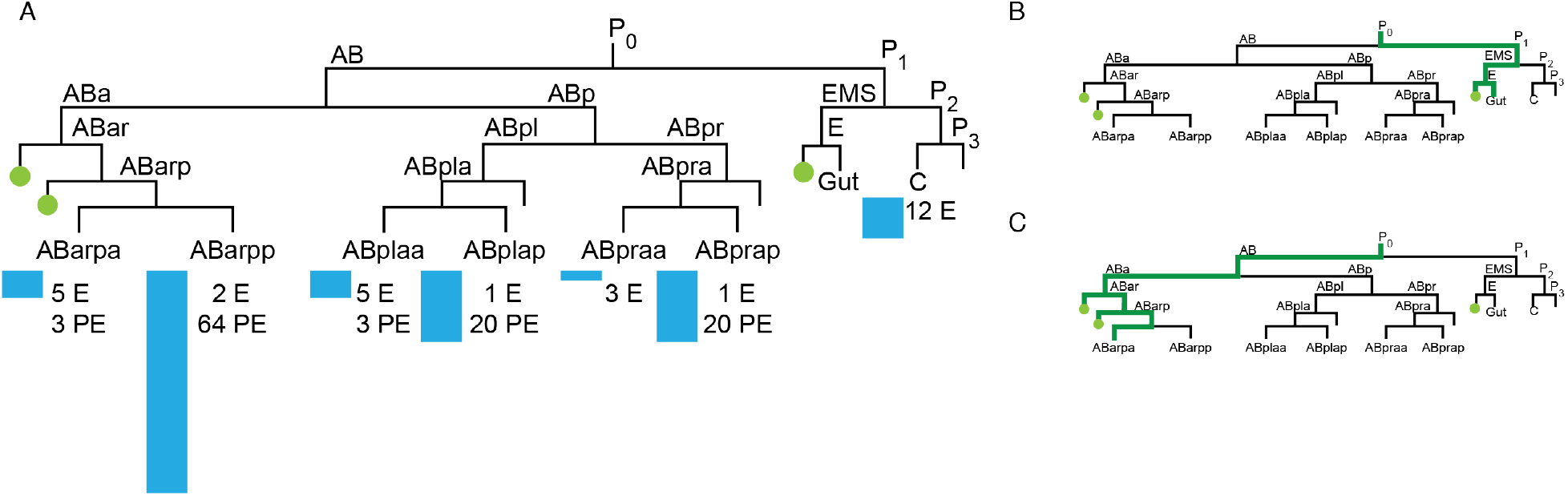
Lineal relatedness of cells that fuse to form syncytial hypodermal cell hyp7. A) Lineage tree showing the cell divisions (vertical lines) leading to the seven early blastomeres that give rise to embryonic (E) or postembryonic (PE) cells that fuse with hyp7. The blue bar show the relative contributions to hyp7 and the green circles show the origins of pharyngeal muscle cells that express *sid-4::gfp*. B, C) Examples of extrachromosomal array transmission patterns that could generate the patterns observed in *bli-1* (RNAi) resistant *sid-4::gfp* positive mosaic animals (Figure 6). B) At the first division the array segregates to P1 but not to AB and at the second division segregates to EMS but not to P2. The resulting animal expresses *sid-4::gfp* in the intestine and cells of the posterior pharynx, but lacks *sid-4::gfp* in all hyp7 nuclei. C) At the first division the array segregates to AB but not to P1, at the second division segregates to ABa but not to ABp, and at a subsequent division segregates to ABar, ABarp, or ABarpp. The resulting animal expresses *sid-4::gfp* in cells of the anterior pharynx, but lacks *sid-4::gfp* expression in all or most hpy7 nuclei.

**Figure S2.**
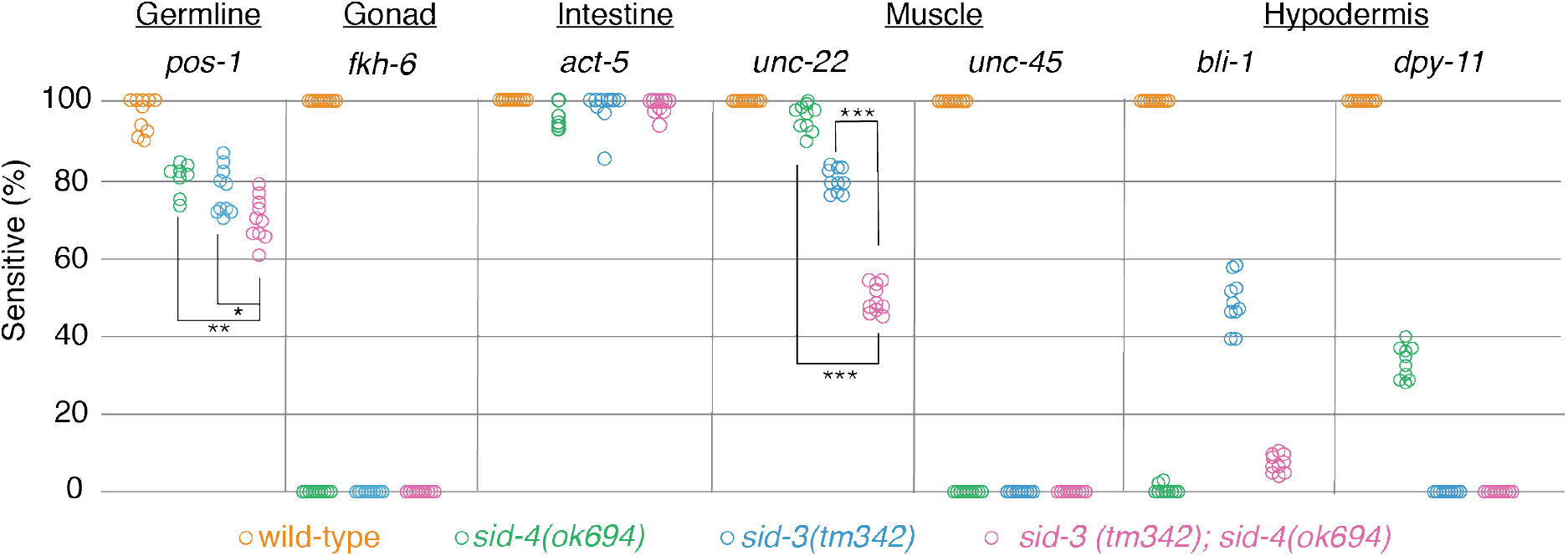
(supporting Figure 7A) RNAi sensitivity of *sid-3* and *sid-4 single* and *sid-3; sid-4* double mutants on a panel of RNAi foods. The F_1_ adult progeny of F_0_ L4 animals placed on RNAi foods were scored for RNAi sensitivity (fraction sensitive on each plate). Each circle represents the plate mean sensitivity for each F_0_ (n=10) with ≥40 F_1_’s per F_0_. P-values (t-test). * <0.01, ** <0.0005, *** <2.3×10^−15^.

**Figure S3.**
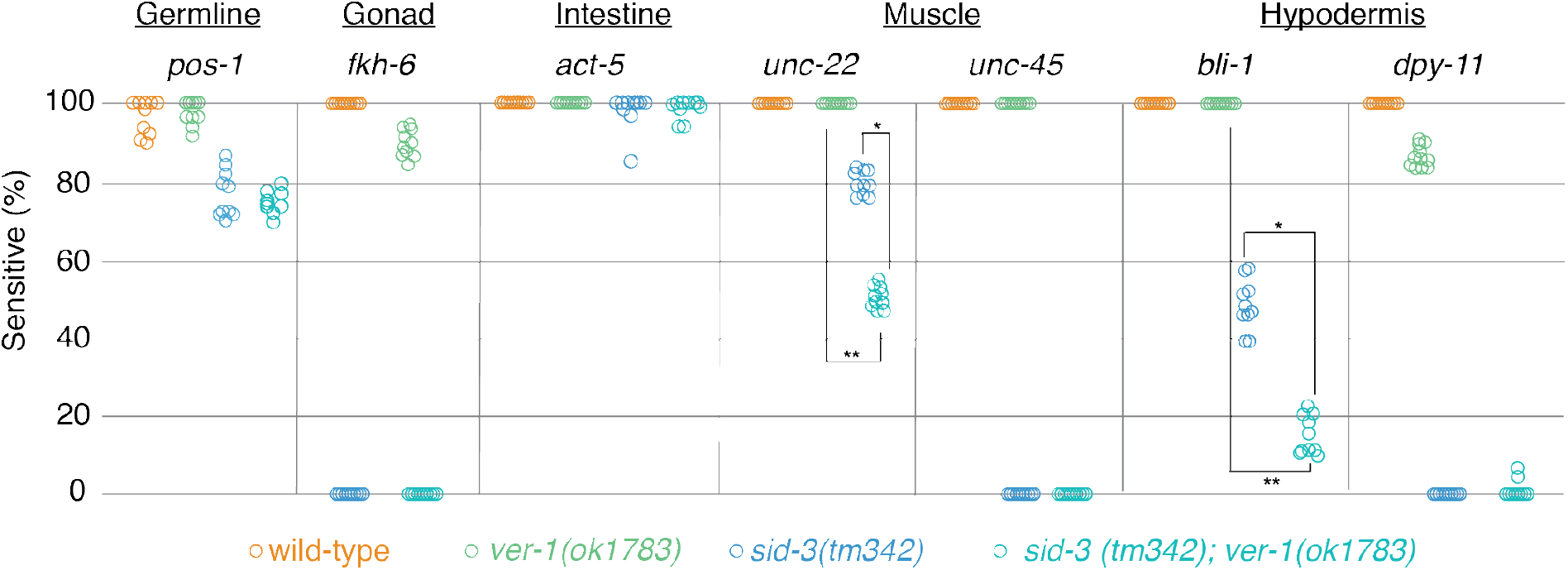
(supporting Figure 7B) RNAi sensitivity of *sid-3* and *ver-1 single* and *sid-3; ver-1* double mutants on a panel of RNAi foods. The F_1_ adult progeny of F_0_ L4 animals placed on RNAi foods were scored for RNAi sensitivity (fraction sensitive on each plate). For *fkh-6*, the proportion of F_1_ adults that contained embryos was scored. Each circle represents the plate mean sensitivity for each F_0_ (n=10) with ≥40 F_1_’s per F_0_. P-values (t-test). * <2.7×10^−11^, ** <1.6×10^−21^.

